# On the organization of the locomotor CPG: insights from split-belt locomotion and mathematical modeling

**DOI:** 10.1101/2020.07.17.205351

**Authors:** Elizaveta M. Latash, Charly Lecomte, Simon M. Danner, Alain Frigon, Ilya A. Rybak, Yaroslav I. Molkov

## Abstract

Rhythmic limb movements during locomotion are controlled by a central pattern generator (CPG) circuits located in the spinal cord. It is considered that these circuits are composed of individual rhythm generators (RGs) for each limb interacting with each other through multiple commissural and long propriospinal circuits. The organization and operation of each RG are not fully understood, and different competing theories exist about interactions between its flexor and extensor components, as well as about left-right commissural interactions between the RGs. The central idea of circuit organization proposed in this study is that with an increase of excitatory input to each RGs (or an increase in locomotor speed) the rhythmogenic mechanism within the RGs changes from “flexor-driven” rhythmicity to a “classical half-center” mechanism. We test this hypothesis using our experimental data on changes in duration of stance and swing phases in the intact and spinal cats walking on the ground or tied-belt treadmills (symmetric conditions) or split-belt treadmills with different left and right belt speeds (asymmetric conditions). We compare these experimental data with the results of mathematical modeling, in which simulated CPG circuits operate in similar symmetric and asymmetric conditions with matching or differing control drives to the left and right RGs. The obtained results support the proposed concept of state-dependent changes in RG operation and specific commissural interactions between the RGs. The performed simulations and mathematical analysis of model operation under different conditions provide new insights into CPG network organization and limb coordination during locomotion.

**Key Point Summary:**

- Limb movements during locomotion are controlled by neural circuits located within the spinal cord. These circuits include rhythm generators (RGs) controlling each limb interacting through multiple commissural pathways.
- The organization and operation of spinal RGs are not fully understood, and different competing concepts exists. We suggest that the operation of RGs is state-dependent, so that with an increase of external excitation the rhythmogenesis changes from “flexor-driven” oscillations to a “classical half-center” mechanism.
- A mathematical model of spinal circuits representing bilaterally-interacting RGs has been developed based on the above suggestion and used to interpret experimental data from intact and spinal cats walking on the ground or tied-belt treadmills (symmetric conditions) as well as on split-belt treadmills with different left and right belt speeds (asymmetric conditions).
- The performed simulations and mathematical analysis of the model under different conditions provide new insights into operation of spinal circuits and limb coordination during locomotion.

## Introduction

It is commonly accepted that the spinal locomotor central pattern generator (CPG) circuits include separate rhythm generators (RGs) that each control a single limb and interact with each other via multiple commissural and long propriospinal pathways. These connections set up phase relationships between the RGs and thus coordinate limb movements and locomotor gait (Rybak *et al.*, 2015; Danner *et al.*, 2017). Each RG is thought to contain two excitatory neuron populations representing flexor and extensor half-centers connected by reciprocal inhibition, whose activity defines the flexor and extensor phases of limb movements, respectively. According to the *classical half-center concept* (Brown, 1914), switching between the flexor and extensor activity phases (for review see (McCrea & Rybak, 2008; Stuart & Hultborn, 2008)) occurs through a so-called *release* mechanism (Wang & Rinzel, 1992) based on an adapting (decrementing) activity of each half-center and mutual inhibition between them. This mechanism does not necessarily require the ability of each half-center to intrinsically generate rhythmic activity, and the resultant RG pattern is usually flexor-extensor balanced, so that the durations of both phases are approximately equal.

The other potential mechanism is based on the intrinsic ability of one or both half-centers to generate rhythmic bursting (Wang & Rinzel, 1992; Skinner *et al.*, 1994; Marder & Calabrese, 1996; Marder & Bucher, 2001). Optogenetic studies in the isolated spinal cord have demonstrated that rhythmic flexor and extensor activities can be evoked in certain conditions independent of each other (Hägglund *et al.*, 2013), confirming that both flexor and extensor half-centers are *conditional* intrinsic oscillators, i.e. capable of endogenous generation of rhythmic bursting activity. Pearson and Duysens have previously proposed a *flexor-driven* concept (so called *swing generator model*, (Pearson & Duysens, 1976; Duysens, 2006); for review see (Duysens *et al.*, 2013)), in which only the flexor half-center is intrinsically rhythmic, hence representing a true RG, while the extensor half-center shows sustained activity if uncoupled and exhibits anti-phase oscillations due to rhythmic inhibition from the flexor half-center.

To meet both concepts, we previously suggested that both half-centers are conditional oscillators, whose ability to intrinsically generate rhythmic bursting depends on the level of excitation (Shevtsova *et al.*, 2015; Danner *et al.*, 2016; Shevtsova & Rybak, 2016; Danner *et al.*, 2017). In this case, a relatively strong excitation of the extensor half-center keeps it in the mode of sustained activity (if uncoupled), whereas a relatively weak excitatory drive to the flexor half-center allows generation of intrinsic oscillations. Therefore, the mechanism for rhythm generation in the RG may vary and, depending on external drives to its half-centers or their level of excitation, it can operate according to the classical half-center or the flexor-driven scenario as was previously demonstrated and analyzed by Ausborn *et al.* (2018).

In the present study, we extend the RG model of Ausborn *et al.* (2018) by assuming that increased activation of the flexor half-center is accompanied by a corresponding decrease in the activity of the extensor half-center. To implement this assumption in the model, we suggested that the external excitatory drive to the flexor half-center simultaneously provides inhibition to the extensor half-center, thus reducing the level of its excitation and directing its operation toward intrinsic rhythmicity. To this end, with an increase of the drive to the RG (with the corresponding increase of oscillation frequency) the operating rhythmogenic mechanism changes from the flexor-driven rhythmicity to classical half-center oscillations with a quasi-balanced flexor-extensor pattern.

To study the behaviors of the proposed RGs in the context of left-right interactions and limb coordination, we incorporated these new RG implementations in the model of left-right circuit interactions in the spinal cord previously described by (Danner *et al.*, 2019). The resultant model included two (left and right) RGs interacting via several commissural pathways presumably mediated by genetically identified V0v, V0d and V3 interneurons. The main goal of this study was to investigate left-right interactions and coordination under different symmetric and asymmetric conditions, which were defined by the same or different drives to left and right RGs, respectively. We assumed these conditions to be, at first approximation, similar to overground or regular tied-belt treadmill locomotion in cats (symmetric conditions) and their stepping on split-belt treadmills with different speeds of the left and right belts (asymmetric conditions). The experimental data were collected from intact and spinal cats in previously published (Frigon *et al.*, 2015; Frigon *et al.*, 2017; Kuczynski *et al.*, 2017) and new experiments. These experimental data were compared with the results of our simulations, in which the modelled circuits operated in similar symmetric and asymmetric conditions. We used these comparisons to evaluate the plausibility of our model and, thus, to formulate important insights into the organization of spinal CPG circuits and their role in limb coordination during locomotion.

## Methods

### Experimental studies

#### Ethical approval

All procedures were approved by the Animal Care Committee of the Université de Sherbrooke in accordance with policies and directives of the Canadian Council on Animal Care (Protocol 442-18). The current dataset was obtained from 11 adult cats (7 females and 4 males) weighing between 3.5 and 5.0 kg. However, only 1 new cat contributed new data during overground locomotion, as data from previous studies were reanalyzed (Frigon *et al.*, 2015) or reused (Frigon *et al.*, 2017) for illustrative or modeling purposes. Before and after experiments, cats were housed and fed in a dedicated room within the animal care facility of the Faculty of Medicine and Health Sciences at the Université de Sherbrooke. As part of our effort to maximize the scientific output of each animal, 10 of 11 animals were used in other studies to answer different scientific questions (Frigon *et al.*, 2013; Thibaudier *et al.*, 2013; D’Angelo *et al.*, 2014; Frigon *et al.*, 2014; Thibaudier & Frigon, 2014; Dambreville *et al.*, 2015; Frigon *et al.*, 2015; Hurteau *et al.*, 2015; Dambreville *et al.*, 2016; Frigon *et al.*, 2017; Hurteau *et al.*, 2017; Kuczynski *et al.*, 2017; Thibaudier *et al.*, 2017; Harnie *et al.*, 2018; Hurteau & Frigon, 2018; Hurteau *et al.*, 2018; Desrochers *et al.*, 2019). The experimental studies complied with the ARRIVE guidelines (Kilkenny *et al.*, 2010) and principles of animal research established by the Journal of Physiology (Grundy, 2015).

#### Surgical procedures

Surgical procedures were described in detail in (Frigon *et al.*, 2015; Frigon *et al.*, 2017) and also apply to the new cat used here. Briefly, we performed all surgical procedures in an operating room with sterilized equipment. Before surgery, the cat was sedated with an intramuscular (i.m) injection of Butorphanol (0.4 mg/kg), Acepromazine (0.1 mg/kg), and Glycopyrrolate (0.01 mg/kg). Induction was done with Ketamine/Diazepam (0.11 ml/kg in a 1:1 ratio, i.m.). The fur overlying the back, stomach, and hindlimbs was shaved. The cat was then anesthetized with isoflurane (1.5 - 3%) using a mask for a minimum of 5 min and then intubated with a flexible endotracheal tube. We confirmed isoflurane concentration during surgery by monitoring cardiac and respiratory rates, by applying pressure to the paw to detect limb withdrawal, and by assessing muscle tone. A rectal thermometer was used to monitor body temperature and keep it between 35°-37°C using a water-filled heating pad placed under the animal and an infrared lamp positioned ~50 cm above the cat. During each surgery, we injected an antibiotic (Convenia, 0.1 ml/kg) subcutaneously and a transdermal fentanyl patch (25 mcg/hr) was taped to the back of the animal 2-3 cm rostral to the base of the tail. During surgery and approximately seven hours later, another analgesic (Buprenorphine 0.01 mg/kg) was administered subcutaneously. After surgery, cats were placed in an incubator and closely monitored until they regained consciousness. At the conclusion of the experiments, cats received a lethal dose of pentobarbital through the left or right cephalic vein.

##### Spinal transection

The spinal cord was completely transected at low thoracic levels in six cats (4 females, 2 males); see (Frigon *et al.*, 2017). A small laminectomy was performed between the junction of the 12th and 13th vertebrae. After exposing the spinal cord, lidocaine (Xylocaine, 2%) was applied topically and injected within the spinal cord. The spinal cord was then transected with surgical scissors. Haemostatic material (Spongostan) was then inserted within the gap and muscles and skin were sewn back to close the opening in anatomic layers. Following spinalization and for the remainder of the study, the bladder was manually expressed 1–2 times each day. The hindlimbs were frequently cleaned by placing the lower half of the body in a warm soapy bath. For training the recovery of hindlimb locomotion see (Frigon *et al.*, 2017).

##### Implantation

All 11 cats were implanted with electrodes to chronically record muscle activity (EMG, electromyography). Pairs of Teflon insulated multistrain fine wires (AS633; Cooner wire, Chatsworth, CA) were directed subcutaneously from 1-2 head-mounted 34-pin connectors (Omnetics Connector Corporation, Minneapolis, MN) and sewn into the belly of selected hindlimb muscles for bipolar recordings. We verified electrode placement by electrically stimulating each muscle through the appropriate head connector channel.

##### Experimental paradigms

Experiments in the 10 cats from previous studies (Frigon et al. 2015, 2017) were performed on an animal treadmill with two independently controlled running surfaces 120 cm long and 30 cm wide (Bertec Corporation, Columbus, OH). Cats performed three locomotor paradigms: 1) Tied-belt locomotion from 0.1 m/s (spinal cats) or 0.4 m/s (intact cats) up to 1.0 m/s in 0.1 m/s increments; 2) split-belt locomotion with one side (slow side) stepping at 0.4 m/s and the other side (fast side) stepping from 0.5 m/s to 1.0 m/s in 0.1 m/s increments; 3) split-belt locomotion with the slow side stepping at 0.1 m/s and the fast side stepping from 0.2 m/s to 1.0 m/s in 0.1 m/s increments (spinal cats only). In spinal cats, the forelimbs remained on a stationary platform with a Plexiglas separator placed between hindlimbs. In the cat that contributed new data, we trained the animal to step along an oval-shaped walkway at self-selected speeds. The walkway has 2.07 m straight paths (0.32 m wide) on each side and we only analyzed data during straight path stepping.

##### Data acquisition and analysis

Videos of the left and right sides during overground and treadmill locomotion were captured with two cameras (Basler AcA640-100 gm) at 60 frames per second with a spatial resolution of 640 by 480 pixels. A custom-made Labview program acquired images and synchronized the cameras with the EMG. Videos were analyzed off-line at 60 frames per second using custom-made software. Contact of the paw and its most caudal displacement were determined for both hindlimbs by visual inspection. We defined paw contact as the first frame where the paw made visible contact with the treadmill surface while the most caudal displacement of the limb was the frame with the most caudal displacement of the toe. We measured cycle duration from successive contacts of the same hindpaw while stance duration corresponded to the interval of time from paw contact to the most caudal displacement of the limb. Swing duration was measured as cycle duration minus stance duration. Durations from 6-15 cycles for each limb were averaged for an episode during treadmill locomotion. In one cat, we obtained and analyzed 44 cycles from 10 runs of overground locomotion.

The EMG was pre-amplified (x10, custom-made system), bandpass filtered (30–1000 Hz) and amplified (x100–5000) using a 16-channel amplifier (AM Systems Model 3500, Sequim, WA). EMG data were digitized (2000 Hz) with a National Instruments card (NI 6032E) and acquired with custom-made acquisition software and stored on computer. The EMG data set shown came from recordings in the anterior sartorius (Srt, hip flexor/knee extensor), the vastus lateralis (VL, knee extensor) and and the lateral gastrocnemius (LG, ankle plantarflexor/knee flexor).

## Mathematical Modeling

We implemented a reduced mathematical model based on the work of (Danner *et al.*, 2017). Simulating flexor and extensor half-centers using activity-based neuron models describing neuron populations (Ermentrout, 1994) significantly simplifies mathematical analysis. The voltage variable of each flexor and extensor units represents the average voltage of the population of flexor and extensor neurons. Such a reduction provides an accurate description of the network dynamics in the CPG circuits controlling mammalian locomotion (Molkov *et al.*, 2015; Ausborn *et al.*, 2018). The CPG network controlling rhythmic locomotion is known to include both excitatory and inhibitory connections between flexor half-centers (Rybak *et al.*, 2013; Molkov *et al.*, 2015; Rybak *et al.*, 2015; Danner *et al.*, 2016; Shevtsova & Rybak, 2016; Danner *et al.*, 2017; Ausborn *et al.*, 2019; Danner *et al.*, 2019). We only included reciprocal inhibition between flexors in the model assuming a net inhibitory interaction. Flexor and extensor half-centers comprising left and right RGs also inhibit each other. Additionally, the model included inhibition from extensors to contralateral flexors. This connection was first introduced by Danner *et al.* (2017) who found that inhibition of flexor half-centers by contralateral extensor stabilize anti-phase left-right alternations in corresponding gaits. In this study we show that this interaction is essential for symmetric leftright alternations and explain the mechanism.

All neurons were modeled using the formalism described in (Rubin *et al.*, 2009) and then used in a number of previous publications (Rubin *et al.*, 2011; Molkov *et al.*, 2014; Molkov *et al.*, 2015; Danner *et al.*, 2016; Molkov *et al.*, 2016; Danner *et al.*, 2017; Ausborn *et al.*, 2018; Ausborn *et al.*, 2019; Danner *et al.*, 2019). Intrinsic bursting properties resulted from slowly inactivating sodium current dynamics. The membrane potential (*V*) of flexors and extensors was governed by the following equation:

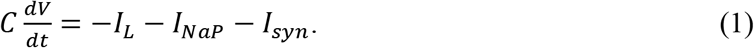

Here, *C* is the capacitance, *t* is time, *I_L_* is the leak current, *I_NaP_* is the slowly inactivating (persistent) sodium current, and *I_syn_* is the synaptic current that is the sum of input currents from other neurons and the excitatory drive current. The leak current and the persistent sodium current were defined in the same manner in flexors and in extensors.

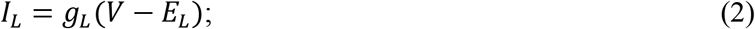

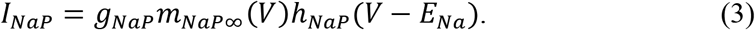

In the expression for the leak current (2), *g_L_* is the conductance of the leak current and *E_L_* is the leak reversal potential. In the expression for the persistent sodium current (3), *g_NaP_* is the persistent sodium maximal conductance and *E_Na_* is the sodium reversal potential. *m*_*NaP*∞_(*V*) is the voltage-dependent steady-state activation function of the persistent sodium current. Persistent sodium current activation is considered to be instantaneous. *h_NaP_* is the persistent sodium inactivation gating variable. The steady state activation functions for persistent sodium activation and inactivation are given by the following expressions:

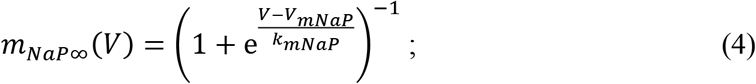

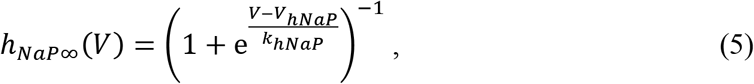

and the dynamics of the persistent sodium inactivation variable were governed by the following differential equation:

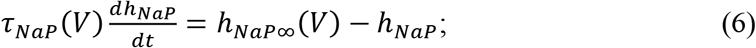

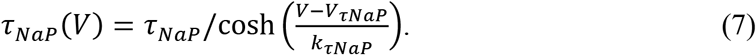

Here, *τ_NaP_*(*V*) is the voltage-dependent time constant for the inactivation of the persistent sodium current. In the gating variable expressions, *V_xNaP_* is the half-(in) activation voltage and *k_xhNaP_* is the (in)activation slope, where *x* ∈ {*m, h, τ*}.

In the differential equation for the membrane potential the third current is the synaptic current *I_syn_* and is defined by the synaptic input from neurons in the network as well as external drives. For flexors, this included input from the contralateral flexor, the ipsilateral extensor, and the contralateral extensor. For extensors, the synaptic current included input from the ipsilateral flexor. In flexors and extensors, drive was implemented as the conductance of an excitatory input. The general expression for the synaptic current in neuron *i* is as follows:

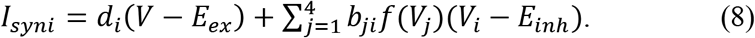

Here, *d_i_* is the excitatory drive to neuron *i* and *V_i_* is the voltage of neuron *i*. *E_ex_* is the reversal potential for the excitatory synaptic currents. *I_syni_* includes the sum over all synaptic inputs from *j* = 1:4 (see Fig. 1). *E_inh_* is the reversal potential for the inhibitory synaptic currents. *b_ji_* is the weight of the synaptic connection from neuron *j* to neuron *i*, which represents the maximal conductance of the corresponding synaptic channel. *f*(*V*) is the activity (normalized firing rate) as a function of voltage and is defined by the following piecewise linear function.

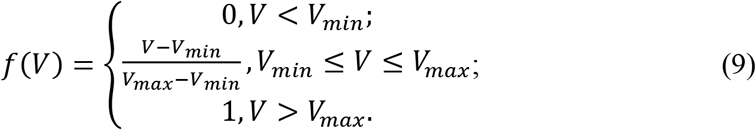

**Figure 1.**
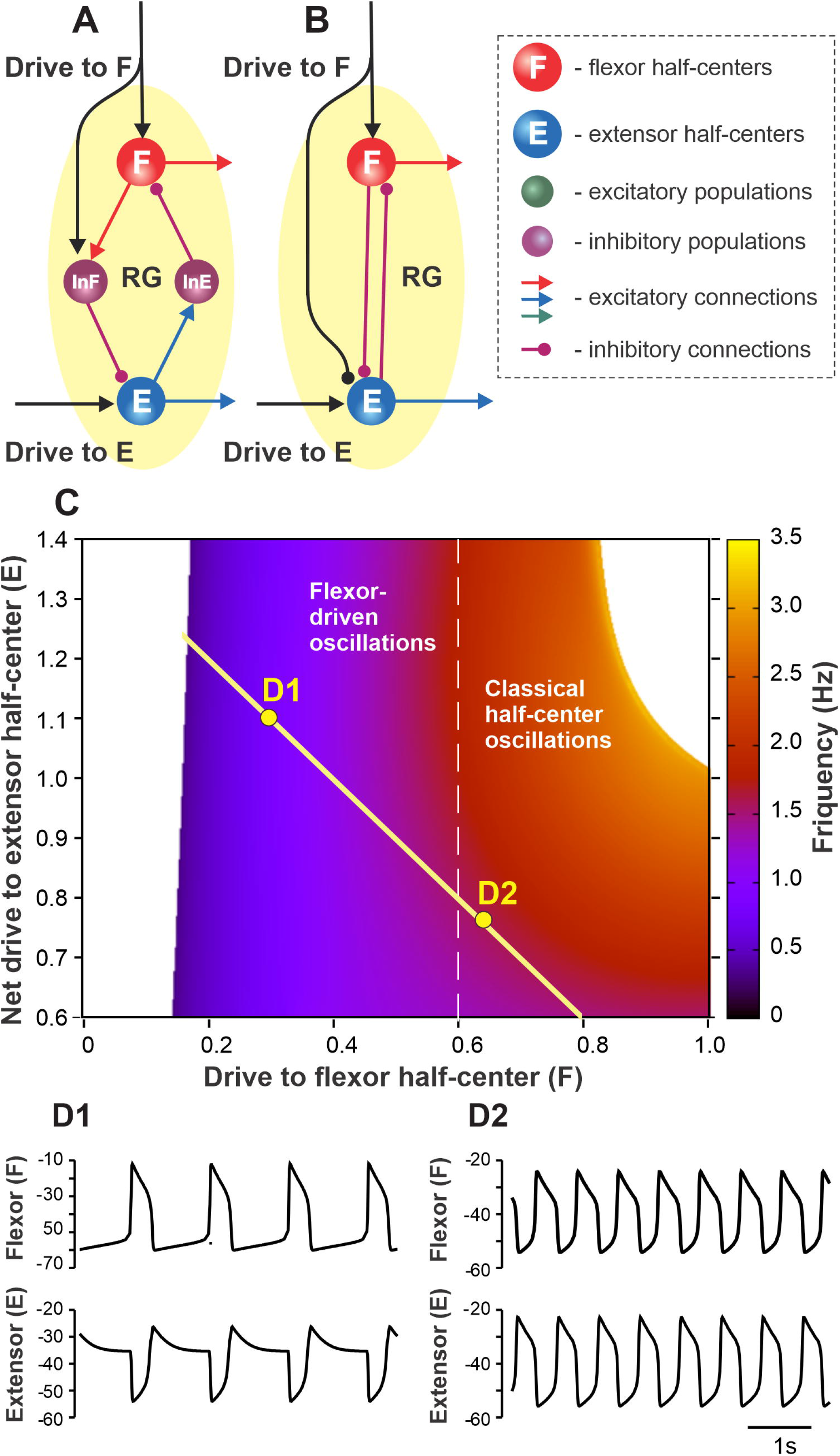
Proposed organization of the single rhythm generator (RG). **A**. Each RG consists of flexor (F) and extensor (E) neural populations (half-centers) inhibiting each other via inhibitory interneuron populations InE and InF. Flexor and Extensor half-centers receive excitatory drives labelled as Drive to F and Drive to E, respectively. Drive to F also excites InF and thus has an inhibitory effect on the extensor half-center. **B**. The simplified model schematic. The inhibitory interneuron pathways are replaced with direct reciprocal inhibition between flexor and extensor half-centers. The net drive to the extensor half-center is defined by the excitatory Drive to E and inhibition from Drive to F. **C**. The dependence of RG bursting frequency on the drive to the flexor half-center and the net drive to the extensor half-center (this representation follows Ausborn *et al.* (2018) methods). A flexor-driven rhythm occurs in the region with relatively high drive to the extensor and low drive to the flexor, i.e. where the flexor half-center is intrinsically rhythmic (to the left from the vertical dashed line). Classical half-center oscillations occur to the right from the vertical dashed line where both flexor and extensor half-centers exhibit tonic activity if decoupled. The hypothetical dependence of the net extensor drive on the flexor drive is shown by yellow line – as the flexor drive increases the net extensor drive decreases due to inhibition from Drive to F to the extensor half-center (see panels A, B). **D1-D2**. Simulated flexor (above) and extensor (below) activity traces for the parameter points labelled as D1 and D2 in panel C.

The activity function *f*(*V*) varies from 0 to 1. Here, *V_min_* and *V_max_* define the voltages at which threshold and saturation are reached, respectively. The values of all parameters are provided in Table 1. In our simulations, the synaptic weights of commissural connections *b*_12_, *b*_21_, *b*_41_ and *b*_32_ were varied, while synaptic weights within each RG *b*_31_, *b*_42_, *b*_13_ and *b*_24_ were fixed.

**Table 1.** Model Parameter Values.

|  |  |
| --- | --- |
| Membrane capacitance (pF) | $C = 20$ |
| Maximal conductance (nS) | $g_L = 2.8, g_{NaP} = 5$ |
| Reversal potentials (mV) | $E_L = -65, E_{Na} = 50, E_{ex} = -10, E_{inh} = -90$ |
| Synaptic weights (nS) | $b_{12} = b_{21}, b_{41} = b_{32}, b_{31} = b_{42} = 0.5, b_{13} = b_{24} = 1$ |
| Threshold and saturation voltage (mV) | $V_{min} = -50, V_{max} = 0$ |
| Time constant (ms) | $\tau_{NaP} = -1500$ |
| $I_{NaP}$ parameters (mV) | $V_{mNaP} = -40, k_{mNaP} = -6, V_{hNaP} = -50,$<br>$k_{hNaP} = 10, V_{\tau NaP} = -100, k_{\tau NaP} = 40$ |

## Results

### Modeling spinal CPG circuits

#### Model of the rhythm generator (RG) controlling a single limb

In the present study, we accepted the model of Ausborn *et al.* (2018) and their suggestion that rhythmic activity in the RG may be based on flexor-driven or classical half-center mechanisms, depending on the level of excitation of flexor and extensor half-centers, both considered conditional bursters. They independently varied flexor and extensor drives and identified parameter areas in which the above mechanisms operate. Here, we extended the model of Ausborn et al. by using the assumption that an increase in activation of the flexor half-center is accompanied by a decrease in the activity of the extensor half-center. Specifically, we assumed that the excitatory drive to the flexor half-center provides inhibition to the extensor half-center (through inhibitory interneurons), reducing the initial level of its excitation (Fig. 1 A, B). In this case, at relatively low drives to the flexor half-center, the frequency of RG oscillations (defined by flexor activity) is low, and the locomotor pattern is not balanced, i.e., has a short flexor and long extensor bursts. An increase in the drive to the flexor half-center increases the RG frequency, making the pattern more flexor-extensor balanced while concurrently reducing the level of excitation of the extensor half-center, shifting the extensor half-center’s operation towards an intrinsically rhythmic state. Figure 1C shows a two-parameter frequency dependence on flexor and extensor drives similar to shown in (Ausborn *et al.*, 2018) that was calculated for a set of parameters used in the present study. According to our suggestion, with the changes in the drive to flexor half-center (Drive to F) and the net drive to extensor half-center (Drive to E minus Drive to F), the parameter point representing a state of RG operation moves along the yellow line intersecting both areas for flexor-driven and classical half-center oscillations (Fig. 1C). Specifically, with an increase of drive to flexor center, the RG operation regimes shifts from the flexor-driven intrinsic oscillations (with short flexor bursts and long extensor bursts, Fig. 1D1) toward the classical half-center mechanism of rhythmicity with a quasi-balanced flexor-extensor pattern (Fig. 1D2).

#### Commissural interactions between RGs controlling left and right limbs

The main goal of this study was to investigate left-right coordination of limb movements under different symmetric and asymmetric conditions. Left-right limb coordination supposedly relies on neural interactions between the two RGs controlling the left and right limbs. The connectome of these interactions was drawn from the model of (Danner *et al.*, 2019). In that model, the left and right RGs interacted via three commissural pathways (Fig. 2A). Two of them, mediated by genetically identified inhibitory V0D and excitatory V0v (V2a-V0v paths, acting via the inhibitory Ini populations) populations of commissural interneurons (CINs), promoted left-right alternation (Talpalar *et al.*, 2013) through mutual inhibition between the left and right flexor half-centers (see also (Shevtsova *et al.*, 2015)). The third pathway, mediated by genetically identified V3 CINs, promoted left-right synchronization via mutual excitation between the left and right extensor half-centers and diagonal inhibition of the contralateral flexor half-centers (Danner *et al.*, 2016; Danner *et al.*, 2019); see Fig. 2A. In the present study, to simplify the model and make it more mathematically tractable, all commissural interactions were replaced by functionally equivalent direct connections, as shown in Fig. 2B.

**Figure 2.**
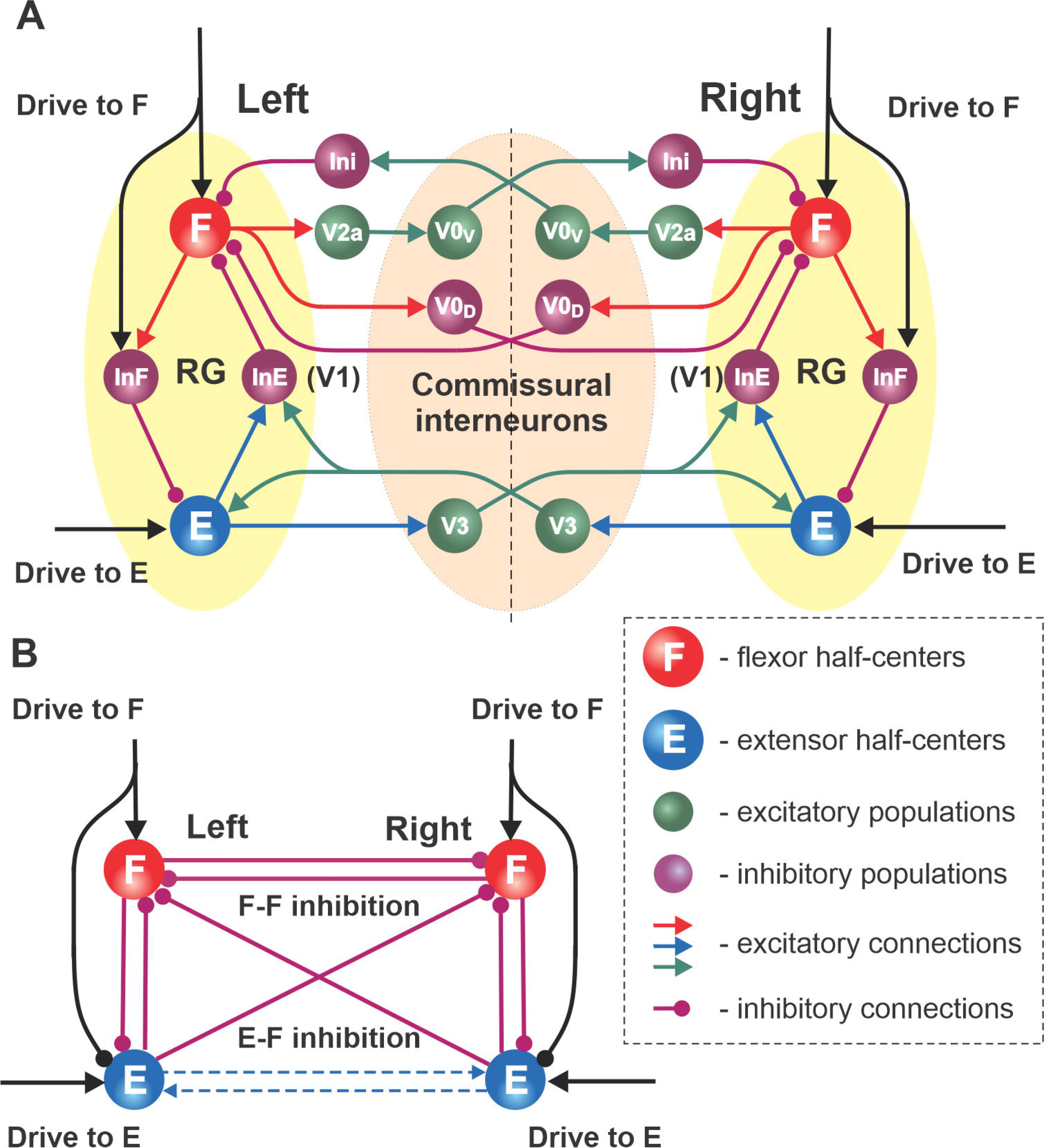
Network interactions between left and right RGs. **A**. Synaptic pathways connecting left and right flexor and extensor interneuron populations proposed by (Danner *et al.*, 2019). The left and right RGs interact through several commissural pathways mediated by different genetically identified commissural interneurons (CINs): V0V, V0D, and V3 types. **B**. Schematic of the simplified model; all CIN-mediated connections are replaced with direct synaptic interactions, i.e. reciprocal inhibition between flexor and extensor half-centers, reciprocal inhibition between flexor half-centers (F-F inhibition) and crossing inhibition from extensor to flexor populations (E-F inhibition). Dashed arrows show the excitatory interactions between extensor half-centers skipped in the simplified model as they are functionally similar to crisscross inhibition.

#### Speed-dependent changes in phase durations during left-right symmetric and asymmetric locomotion

Our objectives was to evaluate the RG circuit organization proposed above by considering their operation in two cases: a symmetric case, when left and right drives vary but remain equal, and an asymmetric case, when one of two drives changes while the other maintains a constant value. We assumed that these two regimes are functionally comparable to regular overground or tied-belt treadmill locomotion (symmetric case) and split-belt treadmill locomotion with different speeds for the left and right belts (asymmetric case). We focused on the analysis of speed-dependent changes in the durations of the main locomotor phases (swing and stance) using data from previous experiments during tied-belt and split-belt treadmill locomotion in intact and spinal cats (Frigon *et al.*, 2015; Frigon *et al.*, 2017) and new experiments performed during overground locomotion in an intact cat.

### Speed-dependent changes in phase durations during left-right symmetric locomotion

#### Left-right symmetric locomotion in cats

Figure 3 shows changes in the cycle duration and durations of swing and stance phases (panel A) and raw activity of representative flexor (Srt) and extensor (LG) muscles (panels B1-B3) during overground locomotion at different self-selected speeds in a freely stepping intact cat. Panels C and D show cycle and phase durations in a group of intact and spinal cats, respectively, during tied-belt treadmill locomotion. In all of these cases, an increase in speed was accompanied by a substantial reduction of stance phase duration with small or absent changes in swing phase duration, consistent with previous studies in cats (Halbertsma, 1983; Frigon & Gossard, 2009; Frigon *et al.*, 2013; Frigon *et al.*, 2014; Frigon *et al.*, 2017). An interesting difference between the three cases shown in Fig. 3 is that during overground locomotion in intact cats, at a speed of ~1.1 m/s, the swing and stance phase durations become equal and then at higher speeds, stance becomes shorter than swing (Fig. 3A). Despite a similar tendency, stance did not become shorter than swing during tied-belt treadmill locomotion in intact (Fig. 3C) or spinal (Fig. 3D) cats. The treadmill locomotion is not usually performed at speeds greater than 1.0 m/s in intact cats, because of safety concerns, as well as in spinal cats, in which the pattern starts to break down. Nevertheless, spinal cats reached swing-stance equality at about 1.0 m/s (Fig. 3D).

**Figure 3.**
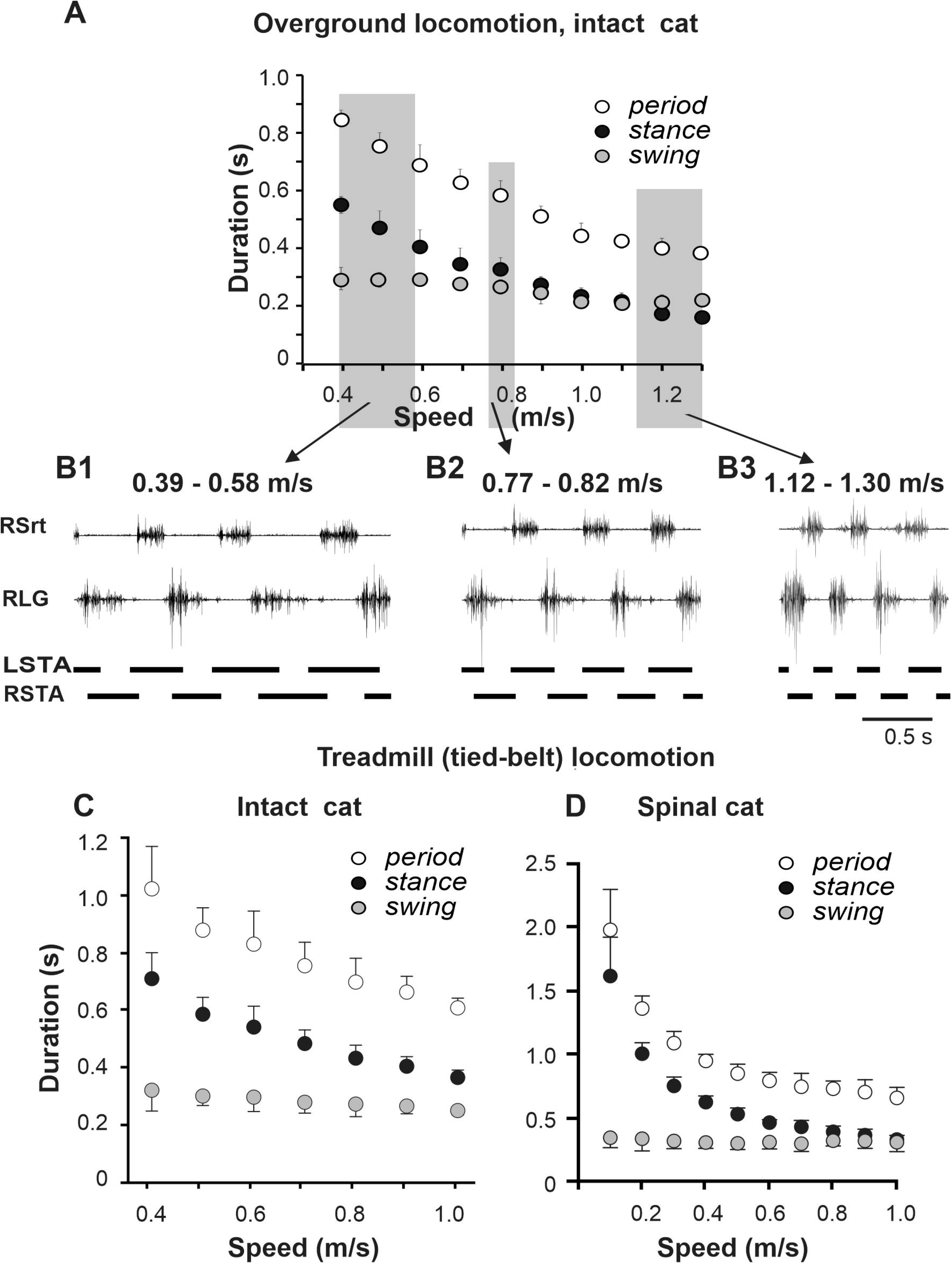
Locomotor cycle and phase durations and muscle activity during overground and tied-belt locomotion across intact and spinal cats. **A.** Cycle and phase durations for the right hindlimb during overground locomotion in an intact cat. The cat stepped in an oval-shaped walkway with 2.07 m straight paths and spontaneously changed speed. We analyzed data from 46 cycles obtained in one session and averaged into 10 bins by rounding to the nearest body speed in 0.1 m/s increments (each data point is the mean ± standard deviation). Note the absence of standard deviations when we only obtained 1 cycle at some speeds. **B1-B3**. Hindlimb muscle activity and phase durations during overground locomotion at 0.39-0.58 m/s, 0.77-0.82 m/s, and 1.12-1.30 m/s in one intact cat. The black horizontal bars at the bottom of each panel show left (LSTA) and right (RSTA) stance phase durations. RLG, right lateral gastrocnemius; RSrt, right sartorius. **C-D.** Cycle and phase durations for the right hindlimb during tied-belt treadmill locomotion in (**C**) intact and (**D**) spinal cats across speeds. We obtained 6-15 cycles in 7 intact and 6 spinal cats and averaged cycle and phase durations for each cat. Each data point is the mean ± standard deviation for the group of intact and spinal cats. **C** and **D** are, respectively, modified from (Frigon *et al.*, 2015) Fig. 2B and reproduced from (Frigon *et al.*, 2017) Fig. 2A, with permission.

#### Simulation of left-right symmetric regime with the model

The schematic of our simplified model is shown in Fig. 2B. In this model there are mutual inhibitory interactions between the flexor half-centers, which combine and simplify two inhibitory pathways mediated by V0d and V0v CINs in Fig. 2A. This inhibition is referred as “flexor-flexor” (or F-F) inhibition. In addition, there are also inhibitory pathways from each extensor half-center to the contralateral flexor half-center (Fig. 2A), which are presumably mediated by V3 CINs through inhibitory populations, such as V1 (Danner *et al.*, 2019). The strength of this connection in the present model is referred to as “extensor-flexor” (or E-F) inhibition. We therefore have four control parameters in the model: the drives to both flexor half-centers (which also define the inhibitory inputs to the extensor half-centers; these drives are equal in the symmetrical case) and F-F and E-F inhibitions.

First, we simulated the changes in locomotor phase durations in response to increasing drive to a single RG (Fig. 4). The external drive to the RGs was increased from 0.2 to 0.8 producing progressively shorter extension at relatively constant flexion duration. With an increase of external drive, the frequency of oscillations increased from about 0.4 to about 1.4 Hz. The increase in frequency (decrease in the period of oscillations) occurred mainly by shortening the extensor phase with minor changes in the duration of the flexor phase. The predominant decrease in extensor phase qualitatively corresponds to the change in the duration of stance and swing phases observed with increasing locomotor speed in experimental studies (Fig 3A). Note that in our simulations, the flexor and extensor phases become equal at drive values of about 0.7, after which extension becomes shorter than flexion (similar to that in Fig. 3A). This reversal in flexor-extensor durations occurs in our model because flexor and extensor half-centers receive the same external excitation at a drive value of approximately 0.7 (see Fig. 1C).

**Figure 4.**
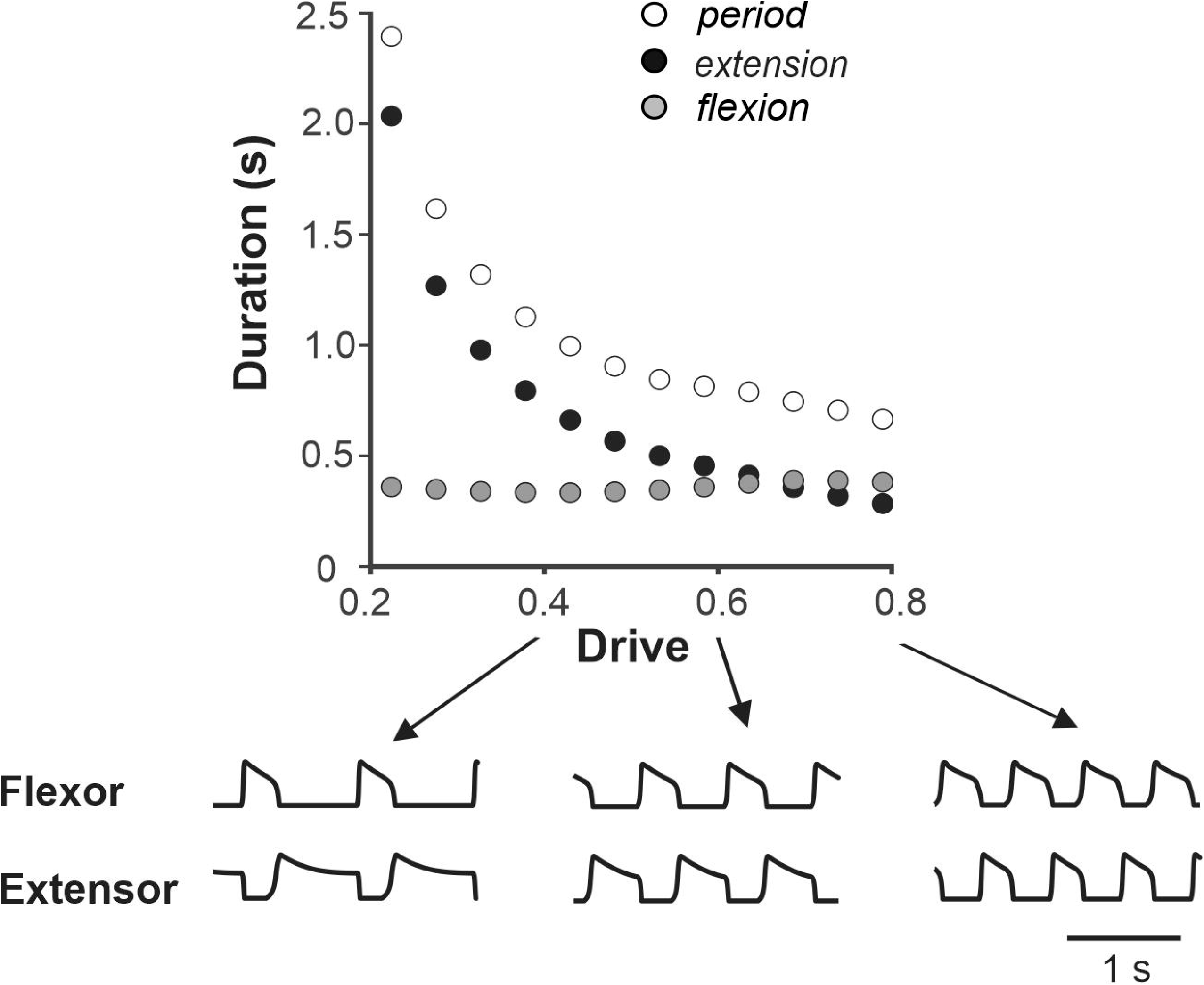
Dependence of the period, flexion and extension on drive to flexor in the model of single RG. Simulations show decreasing duration of extension and relatively constant flexion with increasing drive similar to that during overground tied-belt locomotion in cats with increasing locomotor speed (Fig. 3). Below, exemplar activity traces of flexor and extensor half-centers are shown for low (0.4), medium (0.6) and high (0.8) drive to flexor values.

To explore the system’s behavior in terms of left-right coordination, we simulated the model and identified synchronization patterns while varying inhibition strengths at different drive values. Figure 5A-D shows the parameter plane partitions for four representative drive values corresponding to low and high frequencies. Qualitatively, the F-F inhibition promotes alternating (anti-phase) flexor activity while the E-F inhibition contributes to synchronizing (in-phase) the flexor half-centers due to a phasic reduction in inhibition of flexors during contralateral flexion. Therefore, it is reasonable to expect that at high F-F inhibition and low E-F inhibition (an upperleft corner on Fig. 5 diagrams), the left and right RGs exhibit alternating activity, and at low F-F inhibition and high E-F inhibition their activity synchronizes at all frequencies. These regimes of exact anti-phase and in-phase oscillations are observed in the white and black parameter regions, respectively. There is an overlap between the two regions (shown in grey), which corresponds to bistability in the system, where both regimes can operate depending on the initial conditions chosen. A transition from in-phase to anti-phase oscillations occurs at the boundary between the grey and white regions, which is invariant to the drive (Fig. 5A-D). An opposite transition occurs at the grey-black boundary, which moves up in terms of F-F inhibition with the drive, thus reducing the bistability area.

**Figure 5.**
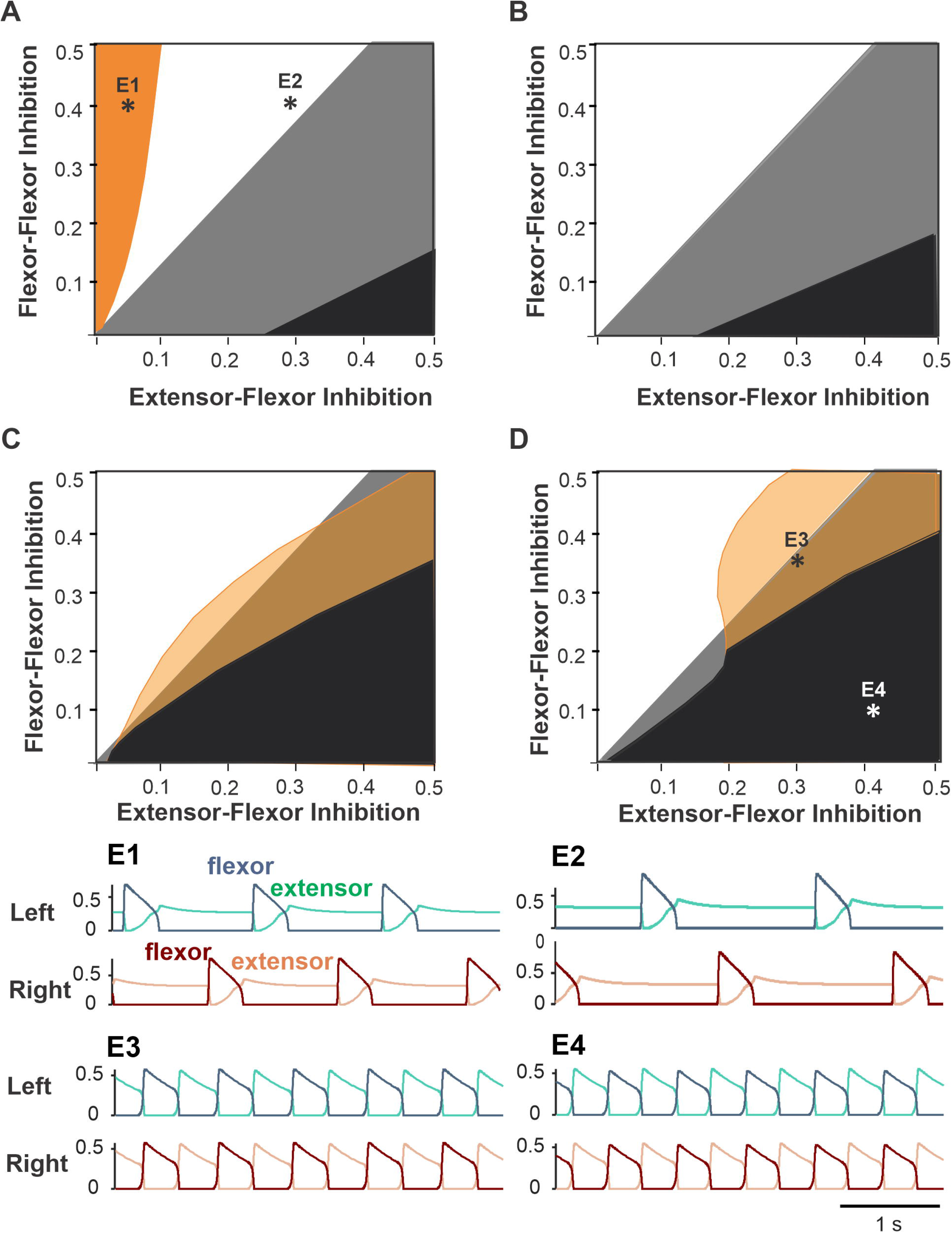
Partitioning of the parameter plane for different coordination patterns. The areas of regimes with different phase relationship between activities of left and right flexor half-centers are shown for varying flexor-flexor (F-F) inhibition and varying crossing extensor-flexor (E-F) inhibition at four different flexor drive values equal to left and right sides (symmetric case). **A**. Drive = 0.3. Orange region: asymmetric alternations of left and right flexor activity – see example activity traces in panel **E1**. The white region corresponds to exact anti-phase left-right alternations (see panel **E2** for an example. The black region corresponds to in-phase left-right synchronization like in panel **E4**. Bistability occurs in the gray region as antiphase and in-phase regimes coexist and can be realized depending on initial conditions. **B**. Drive = 0.4. As we increase drive, the orange region disappears, and the black region of in-phase synchronization grows in size. **C**. Drive = 0.5. With relatively high drive to flexors a new region appears (shown by transparent orange) with small phase difference between flexors (see panel **E3** for an example). The black region of in-phase synchronization increases further. **D**. Drive = 0.65. **E1-E4.** Activity traces of left flexors (blue) and extensors (green) above and the right flexors (dark brown) and extensors (light brown) below corresponding to parameter points labeled accordingly in panels A and D. **E1**. Drive = 0.3, F-F inhibition = 0.4, E-F inhibition = 0.05. **E2.** Drive = 0.3, F-F inhibition = 0.4, E-F inhibition = 0.3. **E3.** Drive = 0.65, F-F inhibition = 0.1, E-F inhibition = 0.4. **E4**. Drive = 0.65, F-F inhibition = 0.33, E-F inhibition = 0.3.

There are also regimes of asymmetric alternations at relatively low (Fig. 5A, orange region) and high (Fig. 5C, D yellow region) drive values corresponding to low or high locomotor frequencies. At low frequencies (i.e. low drive values), this regime is observed at low values of E-F inhibition; it results from post-inhibitory rebound activation of the flexor oscillator after the contralateral flexor deactivates. Slightly higher E-F inhibition strength prevents this post-inhibitory rebound by suppressing the contralateral flexor half-centers for the duration of strong extensor activity in the beginning of the extensor burst. At high locomotor frequencies, the asymmetric alternation regime is practically indistinguishable from pure anti-phase oscillations because the duty cycle is very close to 1/2.

Based on the analysis above, we found that the considered circuit produces robust antiphase alternations of flexor activity in a certain parameter region for all locomotor frequencies. We chose the exemplary point (0.2, 0.4) that belongs to this region for subsequent simulations. However, this particular choice did not make a qualitative difference in the system’s behavior as long as the parameter point chosen belonged to the region of monostable anti-phase oscillations.

### Speed-dependent changes in phase durations and synchronization patterns during left-right asymmetric locomotion

#### Left-right asymmetric locomotion in cats stepping on split-belt treadmills

The split-belt treadmill locomotion experiments, in which animals step on belts with different speeds for the left and right sides, is a common way to study limb coordination during locomotion in cats and humans. Many previous studies in cats demonstrated that both intact and spinal animals adapt to such stepping conditions and demonstrate stable locomotion (Kulagin & Shik, 1970; Forssberg *et al.*, 1980; Frigon *et al.*, 2013; D’Angelo *et al.*, 2014; Frigon *et al.*, 2015; Frigon *et al.*, 2017; Kuczynski *et al.*, 2017). In these studies, we can separate cat locomotion on the split-belt treadmill in two qualitatively different types of conditions: *simple* and *extreme* (Frigon *et al.*, 2017; Kuczynski *et al.*, 2017). In the simple condition, characterized by a relatively small speed difference between moving belts, animals maintain a 1:1 ratio between the number of steps made by left and right limbs. In *extreme* conditions, the animal starts taking more steps on the fast side compared to the slow side resulting in step ratios of 1:2, 1:3, 1:4, etc. (Forssberg *et al.*, 1980; Frigon *et al.*, 2015; Frigon *et al.*, 2017).

The changes in locomotor phase durations during split-belt locomotion of intact and spinal cats in *simple* conditions are shown in Fig. 6; see also (Frigon *et al.*, 2015; Frigon *et al.*, 2017). In both cases, the slow hindlimb (SHL) stepped at a constant speed of 0.4 m/s, whereas the speed of the fast hindlimb (FHL) belt increased from 0.5 to 1.0 m/s. In these conditions, the important characteristics of locomotion observed are (see also (Frigon *et al.*, 2015; Frigon *et al.*, 2017)): (1) The step cycle period remains equal in both hindlimbs (FHL and SHL). (2) In the SHL, the durations of swing and stance phases do not change much. (3) In the FHL, the duration of stance decreases, whereas the duration of swing increases allowing step cycle duration to remain relatively unchanged despite an increase in speed of the FHL. At a FHL speed of ~0.8 m/s, the durations of swing and stance phases become approximately equal and then the swing phase becomes longer than the stance phase at faster FHL speeds (Fig. 6B, right).

**Figure 6.**
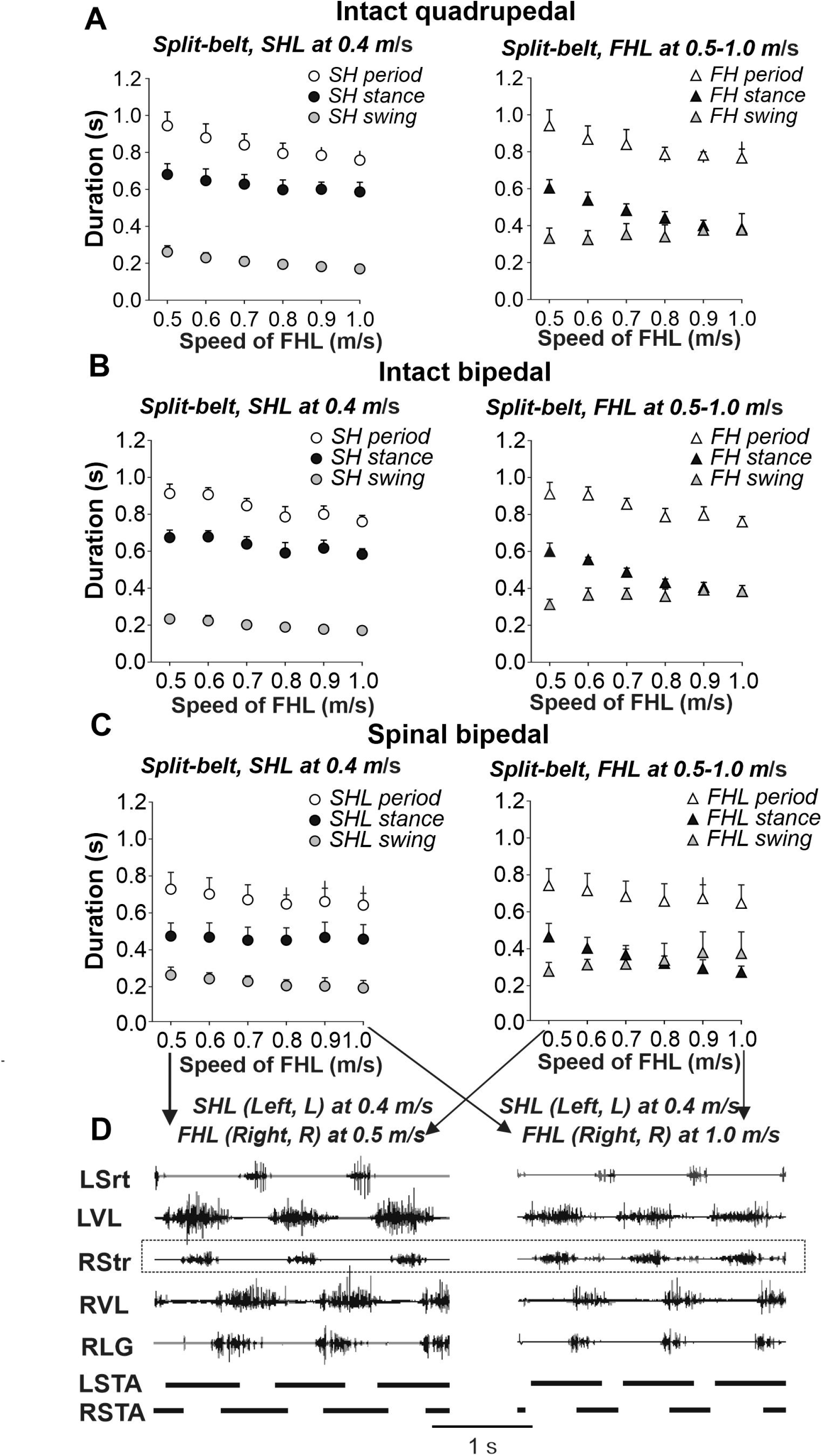
Cycle and phase durations and muscle activity during split-belt locomotion across intact and spinal cats. Cycle and phase durations in (**A-B**) intact and (**C**) spinal cats when the slow hindlimb (SHL) was stepping at 0.4 m/s while the fast hindlimb (FHL) stepped from 0.5 to 1.0 m/s in 0.1 m/s increments. Cycle and phase durations are shown for SHL (left panel) and FHL (right panel). We obtained 6-15 cycles in 5 intact and 6 spinal cats and averaged cycle and phase durations for each cat. Each data point is the mean ± standard deviation for the group of intact and spinal cats. **D**. Hindlimb muscle activity and phase durations during split-belt locomotion with the slow limb stepping at 0.4 m/s and the right hindlimb stepping at 0.5 m/s (left panel) and 1.0 m/s (right panel) in one spinal cat. The black horizontal bars at the bottom of each panel show left (LSTA) and right (RSTA) stance phase durations. Data shown are from cat BL (Frigon *et al.*, 2017). L, left; R, right; LG, lateral gastrocnemius; Srt, sartorius; VL, vastus lateralis. **A, B** are recalculated based on data shown in Figs. 2D, F in (Frigon *et al.*, 2015), **C** is reproduced from (Frigon *et al.*, 2017) Figs. 5A, B, and **D** is reproduced from (Frigon *et al.*, 2017) Figs. 4A, B with permission.

The locomotor characteristics of intact and spinal cats differ in *extreme* conditions, when the speed ratio between the slow and fast belts are set to 1:3 and more, up to 1:10 (Frigon *et al.*, 2017; Kuczynski *et al.*, 2017). In this case, the locomotor pattern changes in such a way that cats take more steps on the fast side than on the slow side. Specifically, at 1:3 and 1:4 speed ratios, the limbs on the fast side perform 2-3 steps for every step of the limb on the slow side (1:2 and 1:3 coordination pattern), whereas at ratios of 1:5 or higher, 1:4 and 1:5 coordination pattern were observed (Frigon *et al.;* Kuczynski *et al.*, 2017). Despite inter-animal variability, both intact (Kuczynski *et al.*, 2017) and spinal (Frigon *et al.*, 2017) cats exhibit 1:2+ coordination patterns.

#### Modeling asymmetric CPG operation

To simulate asymmetric conditions corresponding to different speeds of the treadmill belts, we varied drives to the left and right RGs in our model independently (Fig. 2B), so that if disconnected they would produce unsynchronized flexor/extensor alternations with different frequencies. Due to commissural interactions, the model generated different synchronization patterns depending on parameters. We assumed that the left RG receives a smaller drive. This corresponds to a triangular region above the bisector in the bifurcation diagram shown in Fig. 7A. The bisector of the bifurcation diagram corresponds to equal drives, where exact anti-phase left-right alternations of flexor activity are produced at the commissural connection weights chosen.

**Figure 7.**
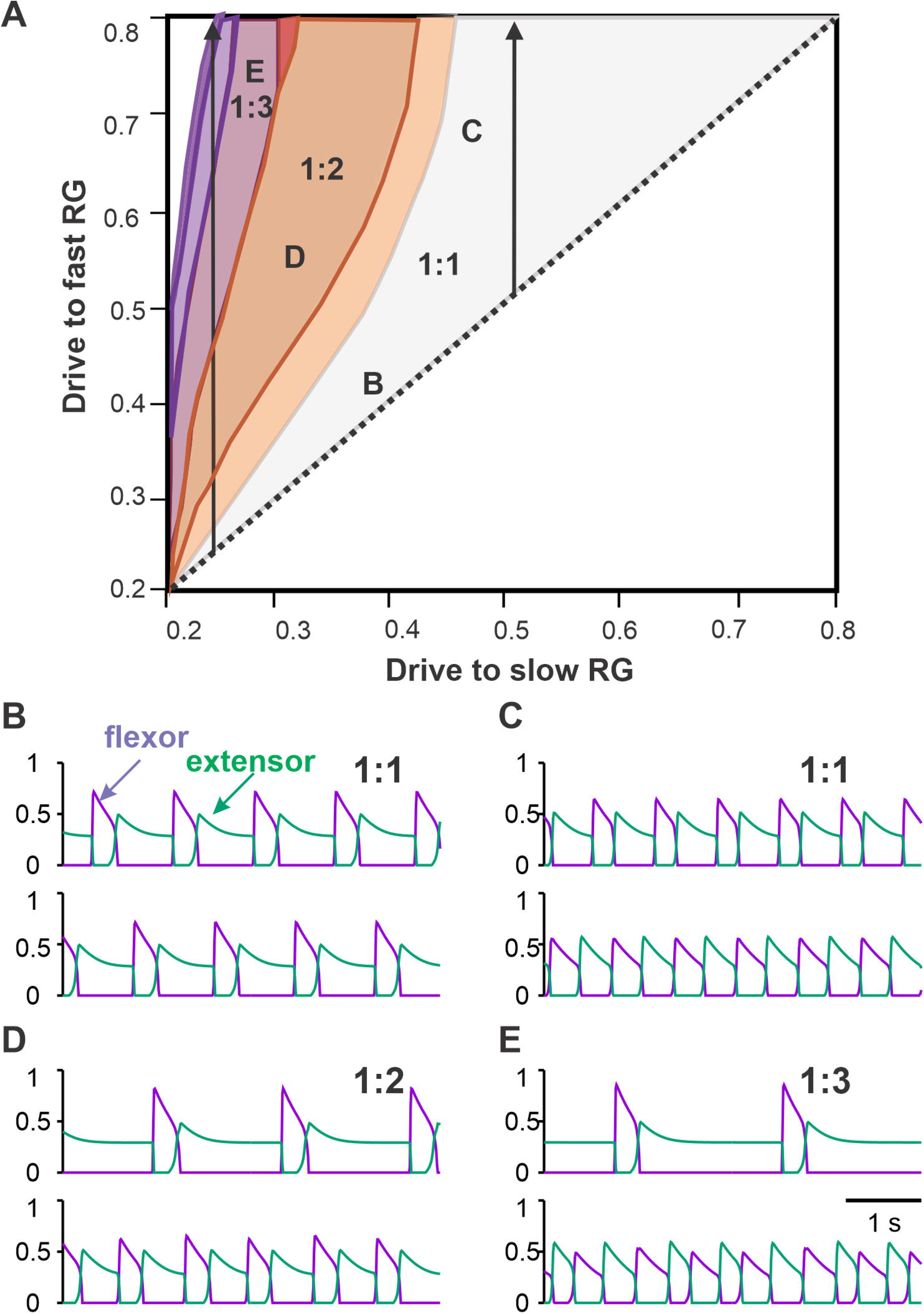
Coordination patterns in the model with asymmetric drives to left and right RGs. **A**. Parameter regions corresponding to different numbers of steps on the fast (right) side per one step on the slow (left) side. The region of a single fast flexor burst for each slow flexor burst is labeled 1:1. Regions of multiple right flexor bursts for each left flexor burst are labeled 1:2, 1:3, etc. The left arrow shows regions of 1:2, 1:3 and higher asymmetric gaits with increasing fast flexor drive at a low strength slow flexor drive, corresponding to extreme experimental conditions. The right arrow shows increasing fast flexor drive and a constant slow flexor drive of moderate strength, corresponding to the simple conditions in split-belt experiments. **B-E**. Examples of activity traces are shown for left (above) and right (below) flexors (violet) and extensors (green) corresponding to the parameter points labelled accordingly in panel A. **B**. As in the tied-belt paradigm, symmetric drive distribution to the left and right flexors produces synchronous antiphase oscillations. **C**. As we increase the drive to the right flexor while keeping the drive to the left flexor at 0.5, the gait becomes asymmetric with longer flexion and shorter extension on the fast right side. **D**. When the drive ratio to right and left flexors is high enough, the right flexors bursts twice for every extensor burst in a 1:2 asymmetric gait. **E**. Even higher drive ratio results in three right flexor bursts for each left flexor burst in a 1:3 asymmetric gait.

As we start changing the drives to the fast RG, both RGs remain synchronized (1:1 region in Fig. 7A), however left and right oscillations become asymmetric. Flexor bursts in the fast RG occur at progressively shorter intervals after flexor bursts. When the drive to the fast RG becomes significantly larger that the drive to the slow RG, the flexor bursts of the fast RG start occurring immediately when flexor bursts of the slow RG end (Fig. 7C). In addition, the duration of the flexor bursts of the fast RG becomes progressively longer (see below in relation to Fig. 8A, B). These behaviors correspond to the *simple* asymmetric conditions, described above, where a 1:1 coordination pattern is maintained.

**Figure 8.**
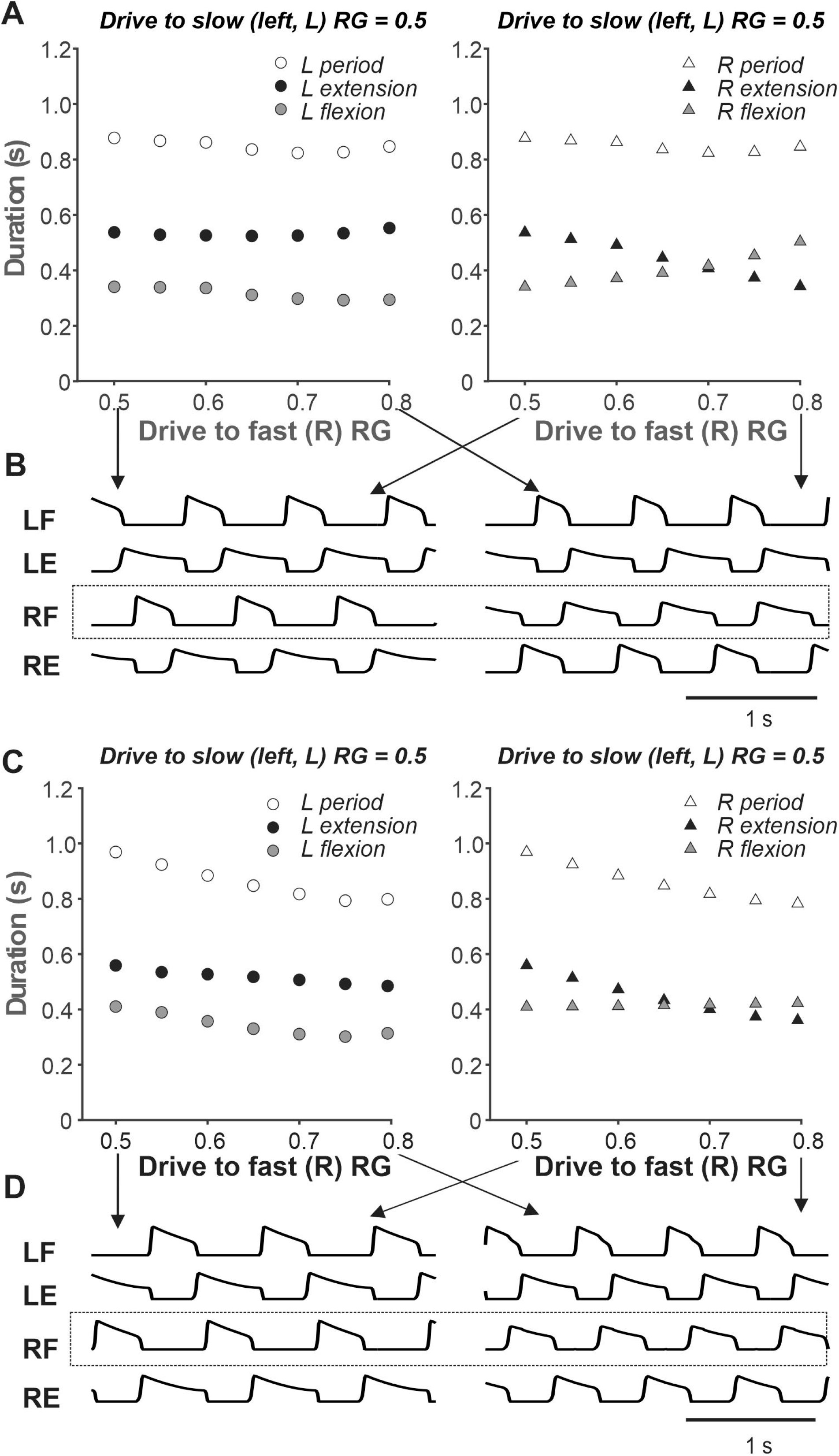
**Simulations of asymmetric CPG activity** as the drive to the slow (left) flexor is kept constant and the drive to the fast (right) flexor is increasing. **A**. The period, flexion and extension duration of the left (slow) and right (fast) RGs as simulated using the model are shown in the left and right panels, respectively. Flexion and extension duration of the slow RG remain fairly constant (left panel). Flexion phase of the fast RG increases in duration while the extension phase of the fast RG shortens in duration (right panel) as in split-belt experiments (see Fig. 6). **B**. Activity traces of flexor and extensor half-centers in symmetric conditions (Drive to both flexors = 0.5, left panel) and asymmetric conditions (Drive to slow flexor = 0.5, Drive to fast flexor = 0.8, right panel). **C-D**. For comparison, same as A-B but with inhibitory effect of the flexor drive on the extensor activity excluded from the model. Drive to both extensor half-centers is kept constant at 0.7. **C.** The period, flexion and extension durations of the slow (left) RG all decrease with increasing drive to the fast (right) RG (left panel). The flexion duration of the fast (right) RG remains constant unlike in split-belt experiments. **D**. Flexor and extensor activity traces of left and right RGs for the minimal (0.5) and maximal (0.8) values of the drive to the fast (right) flexor corresponding to simulations in panel C are shown in left and right panels, respectively. L, left; R, right; RG, rhythm generator; LF, left flexor; RF, right flexor; LE, left extensor; RE, right extensor.

When the frequency of the slow RG is relatively low because of a low drive to the slow RG (left part of the bifurcation diagram in Fig. 7A), a transition to *extreme* conditions (1:2+ coordination patterns) occurs as we increase the drive to the fast RG further (see above). In the 1:2 regime, one flexor burst of the slow RG corresponds to two flexor bursts of the fast RG (1:2 area in Fig. 7A). In this regime, the first flexor burst of the fast RG starts immediately after the flexor burst of the slow RG ends (Fig. 7D). Further increases in the drive to the left (fast) RG leads to the emergence of 1:3+ patterns (Fig. 7E), similar to that observed in *extreme* conditions in intact and spinal cats (see above). Between 1:1 and 1:2 regions, there is an area of intermittent regimes where either one or two flexor bursts can be produced by the fast RG during the extension phase of the slow RG, which is commonly observed experimentally (Frigon *et al.*, 2017).

#### Changes in locomotor phase duration in a simple asymmetric regime (1:1)

Modeling and analysis of locomotor characteristic changes in the simple condition is more functionally relevant than the *extreme* cases because it occurs frequently during everyday locomotion, such as stepping along a circular path or when turning. Also, these changes provide an indirect test for the CPG network organization predicted by the model.

Figure 8A, B shows our simulation of such a *simple* asymmetric case, when the drive to the slow RG was kept constant at 0.5, while the drive to the fast RG increased from 0.5 to 0.8 (see the corresponding arrow in Fig. 7A). Similar to the experimental studies during split-belt locomotion in a simple asymmetric case shown in Fig. 6, despite the left-right asymmetry, the oscillation period remained almost constant and was largely defined by the slow side. Similarly, the durations of flexor and extensor phases were relatively constant on the slow side but changed dramatically on the fast side with increased drive (Fig. 8A, B). The most important feature of the simulated behavior (which corresponded to experimental data in Fig. 6) was the increased duration of flexion in the fast RG occurring with increased drive to that RG. We can qualitatively explain this phenomenon in the model as follows. On the slow side, the flexor half-center of the slow RG operates in a rhythmic mode, while its extensor half-center operates in a regime of tonic activity (if disconnected) as it receives higher excitatory drive. Therefore, the generation of flexor bursts in the slow RG occurs endogenously after a well-defined recovery period, which is almost unaffected by the synaptic inputs it receives from the other side (fast RG). On the fast side, however, once the net drive to the extensor half center is low enough (recall that based on our assumption an increase in drive to the flexor half-center is accompanied by a decrease in drive to the extensor half-center; see above), the extensor half-center goes into an intrinsically rhythmic mode, meaning that the duration of extension and its interburst intervals start to depend on intrinsic burst recovery mechanisms. At the same time, the flexor half-center of the fast RG receives increasingly more excitation, so flexor burst termination becomes more dependent on the onset of extensor half-center inhibition rather than on the flexor’s endogenous deactivation. With a progressive reduction of net drive to the extensor half-center, the recovery period for extensor activity gets longer, which extends flexion duration. Therefore, the phenomenon of increasing duration of flexion in the fast RG results from changing the rhythmogenesis mechanism in the fast RG from an intrinsic generation of flexor oscillations to the classical half-center mechanism that was implemented in our RG model.

To illustrate this further, we removed inhibitory external inputs to both (left and right) extensor half-centers (that provided the above transition in the rhythmogenic properties of the extensor half-centers) and replaced them with a constant excitatory drive of 0.7 (see Fig. 1C). In this case, rhythmogenesis was always based on intrinsic bursting of flexor half-centers without switching to the classical half-center mechanism. The results of these simulations are shown in Fig. 8C, D. Note that (a) the duration of the flexor phase on the fast side never increases, and (b) the step-cycle duration on both sides clearly decreases with increasing drive to the fast RG, both contradicting to experimental observations (see Fig. 6).

## Discussion

### Organization and operation of spinal rhythm generators (RGs) controlling limb movements during locomotion

There are currently two major competing concepts concerning the organization and operation of spinal neuronal RGs. In the classical half-center concept (Brown, 1914), flexor and extensor half-centers do not require intrinsic rhythmic properties (for review see (McCrea & Rybak, 2008); Stuart and Hultborn (2008)). Both half-centers operate in qualitatively similar conditions with phase switching defined by a release mechanism (Wang & Rinzel, 1992) that is based on adapting (decrementing) activity of each half-center and mutual inhibition between them. In the classical half-center, the durations of flexor and extensor phases are balanced (or equal). These durations and the corresponding duty cycles can be easily changed by the level of half-center activation or by external drive. At the same time, the control of RG oscillation frequency in this case is problematic as the oscillation period is not very sensitive to the external drive in half-center oscillators (Daun *et al.*, 2009).

In contrast, with the flexor-driven concept (Pearson & Duysens, 1976; Duysens, 2006), the RG rhythm and pattern is defined by the intrinsically rhythmic flexor half-center, while the extensor half-center has sustained activity if uncoupled and only exhibits rhythmic bursting through rhythmic inhibition from the flexor half-center (for review see Duysens *et al.* (2013)). Thus, the frequency of intrinsically generated flexor bursting explicitly depends on flexor half-center excitation. The distinctive feature of this regime is that the flexor bust duration does not change much and most previously suggested intrinsic oscillatory mechanisms, such as those based on intracellular dynamics of ionic concentrations or slow inactivation of ionic channels (Jasinski *et al.*, 2013; Molkov *et al.*, 2015), produce duty cycles of bursting usually less than 0.5 and are likely to operate at low frequencies with short flexor phases and long extensor bursts.

Both concepts have support in certain conditions. Ausborn *et al.* (2018) demonstrated that both mechanisms can operate depending on the state of half-centers defined by their level of excitation. Here, we used and refined this idea, by suggesting that (a) at low frequencies the extensor half-center is highly excited and operates in a regime of tonic activity, and (b) an increase in excitation of the flexor half-center, which initially operates in the intrinsic bursting regime, is accompanied by a decrease of excitation of the extensor half-center. Mechanistically, such a decrease of the extensor half-center activation may result from a reduction of excitatory afferent inputs to the extensor half-center when unloading the limb at the stance-to-swing transition (Pearson, 1995; Dietz & Duysens, 2000). With concurrent increases in flexor and extensor drives, the RG transitions from a flexor-driven mechanism (when the frequency changes mostly with extension duration while flexion duration remains relatively unchanged) to the classical half-center mechanism (when stepping is controlled by changes in the duty cycle at a relatively constant frequency).

To test this idea, we incorporated the above RGs in a model of spinal CPG circuits with reciprocal commissural interactions and used this bilateral RG model to simulate speed-dependent changes in the locomotor pattern of intact and spinal cats in symmetrical (during overground and tied-belt locomotion) and asymmetrical (during split-belt treadmill locomotion) conditions. The experimental data from previously published (Frigon *et al.*, 2015; Frigon *et al.*, 2017; Kuczynski *et al.*, 2017) and new experiments were analyzed. The model reproduced and explained a series of experimental findings, including (a) the reversal in flexor and extensor phase durations with an increase of locomotor speed during left-right symmetric locomotion, and (b) the maintenance of step cycle period during split-belt locomotion due to adjustment of the flexor duty cycle. The results of these simulations provide strong support for the proposed organization and operation of spinal locomotor circuits.

### Organization of left-right commissural interactions in the spinal cord: the role of V3-mediated commissural pathways

In the present model, the interactions between left and right RGs were based on the model by (Danner *et al.*, 2019). Importantly, that model was derived from experiments on symmetric (bilateral) and asymmetric (unilateral) optogenetic stimulations of commissural V3 neurons involved in left-right coordination performed in the same study. Interestingly, unilateral stimulation produced effects that were qualitatively similar to some features of split-belt locomotion. They provided strong evidence that spinal V3 CINs are involved in left-right limb coordination via two pathways: through mutual excitation between the left and right extensor half centers of the RGs and, importantly, via crossed inhibition from extensor half-centers to contralateral flexor half centers through an additional inhibitory interneuron population (presumably V1) (see. Fig. 2A). In the present study, we show that the commissural inhibition of flexor half-centers by the contralateral extensor half-centers (see Fig. 5 and related texts) is critically important for the stability of anti-phase flexor oscillations at low frequencies in symmetric conditions, which corresponds to a normal locomotor pattern. Therefore, our study provides additional support for the important role of V3 CINs and the existence of inhibitory commissural pathways from extensor half-centers to contralateral flexor half-centers, mediated by V3 and (presumably) V1 interneurons (Danner *et al.*, 2019). Although this prediction still awaits experimental testing, crossed inhibition to flexors (by afferent stimulation) has been observed in anesthetized preparations (Jankowska *et al.*, 2005; Jankowska & Edgley, 2010) and during locomotion in intact cats (Hurteau *et al.*, 2018) as well as in mouse (Laflamme & Akay, 2018) and human (Mrachacz-Kersting *et al.*, 2017) studies.

In summary, our analysis of the model allowed us to evaluate the specific roles of the two types of inhibitory commissural interactions (called here *flexor-flexor* and *extensor-flexor* inhibition) in left-right coordination. The flexor-flexor inhibition, presumably mediated by V0 CINs (Talpalar *et al.*, 2013; Shevtsova *et al.*, 2015), supports left-right alternation and its weakening may stabilize left-right in-phase synchronization. The extensor-flexor inhibition, presumably mediated by V3 CINs and V1 interneurons (Danner *et al.*, 2019), ensures that left and right activities alternate in a strict out-of-phase manner in symmetric conditions.

### Insights from symmetric locomotion

It is well known that during normal locomotion in cats and humans, an increase of speed is accompanied by a significant reduction of stance phase duration with or without a minor reduction of swing phase duration (Grillner *et al.*, 1981; Halbertsma, 1983; Frigon & Gossard, 2009; Frigon *et al.*, 2013; Frigon *et al.*, 2014; Frigon *et al.*, 2017) see also Fig. 3. This observation seems to support the flexor-driven concept of locomotor rhythm generation. However, in intact and spinal cats, increasing locomotor speed produces a more balanced pattern, with stance duration approaching and even becoming shorter than swing duration. This is clearly observed during overground locomotion in intact cats (Fig. 3A). We suggest that when approaching the point of equality between phases, rhythmogenesis shifts towards the classical half-center mechanism. The observation indirectly supporting this view is that after the point of equality, the oscillation period (and hence the frequency) saturates and does not change much, which is a typical feature of classical half-center dynamics (Daun *et al.*, 2009; Ausborn *et al.*, 2018).

### Insights from asymmetric split-belt treadmill locomotion

Previous experimental studies in cats using split-belt treadmill locomotion demonstrated that the mammalian spinal cord has a remarkable adaptive capacity for left–right coordination, from simple to extreme conditions (Forssberg *et al.*, 1980; Halbertsma, 1983; Frigon *et al.*, 2013; Frigon *et al.*, 2015; Frigon *et al.*, 2017). In *simple* conditions, with slow/fast speed ratios of up to 1:2.5 (0.4:1.0 m/s), animals maintain the period of oscillations (and frequency) almost unchanged and compensate for the reduction of stance phase duration on the fast belt by a corresponding increase of the duration of the swing phase (Frigon *et al.*, 2015; Frigon *et al.*, 2017); see Fig. 6. Our model was able to reproduce this feature specifically due to the implementation of our suggestion, that increased activation of the flexor half-center in each RG is accompanied by a reduction in the activity of the corresponding extensor half-center. This implementation leads to a switch in the rhythmogenic mechanism of the fast RG from flexor-driven oscillations to the classical half-center mechanism (Fig. 8A, B). Removing this feature from the model leads to constant swing duration accompanied by a noticeable increase of oscillation frequency in both limbs (RGs) with increasing drive to the flexor half-centers (Fig. 8C, D), contradicting the experimental results, shown in Fig. 6.

Experimental studies of cat locomotion on split-belt treadmills in *extreme conditions*, with slow/fast speed ratios of 1:3 and more (Frigon *et al.*, 2017; Kuczynski *et al.*, 2017) showed that cats use a specific strategy to stabilize locomotion by taking multiple steps on the fast side per step on the slow side. Moreover, although there was some variability between animals, both intact (Kuczynski *et al.*, 2017) and spinal (Frigon *et al.*, 2017) cats exhibit 1:2, 1:3 or 1:4 coordination patterns corresponding to 2, 3 or 4 steps on the fast side per step on the slow side, respectively. To simulate these behaviors, we applied different drives to the left and right RGs in the model, assuming that these conditions are qualitatively similar to the extreme case of split-belt locomotion. Under these conditions, the model predicts that the number of different coordination patterns depends on the value of the drive to the slow RG (Fig. 7). For relatively high drives to the slow RG (>0.45), only a 1:1 coordination pattern is possible, which corresponds to simple conditions in split-belt locomotion (see above). However, if the drive to the slow RG is smaller, 1:2+ coordination patterns become possible. For example, for a slow RG drive value of 0.4, as the drive to the fast RG increases, there is a transition from 1:1 to 1:2 coordination pattern, but no 1:3 regime exists, while for a slow RG drive value of 0.25, as the fast RG drive progressively increases, the system undergoes 1:1, 1:2, 1:3 and 1:4 regimes. Qualitatively similar behavior is observed in extreme split-belt locomotion where in order to achieve higher order coordination patterns, one has to set lower speeds of the slow belt.

### Limitations, functional considerations, and future directions

In this study we show that a relatively simple functional connectome between populations of interneurons providing output to flexor and extensor motoneurons that control a pair of limbs can explain a variety of coordination patterns emerging in split-belt experiments. The mathematical model we developed allowed us to formulate a novel hypothesis about general mechanisms of locomotor phase duration control suggesting that variation of the excitatory drive to the flexor half-centers is accompanied by an opposite change in the drive to the extensor half-centers. However, our model does not provide any specifics on neuronal pathways mediating these interactions.

What would be the benefit of switching from a flexor-driven RG operation to a classic half-center mode with increasing speed? Although we can only speculate, the goal of the spinal locomotor network might be to optimize efficiency or balance (avoid falling). At slow to moderate speeds, the stance duration is long and inputs from group I/II extensor muscle afferents and paw pad cutaneous afferents have a relatively long time to regulate stance duration and adjust/correct for destabilizing perturbations. Thus, at slow to moderate speeds, a flexor-driven RG mode is less costly and more efficient. However, as speed increases, stance duration also decreases and afferent inputs do not have as much time to adjust or correct for postural perturbations. As such, at high speeds, a classic half-center mode, whereby both stance and swing phase durations are balanced, becomes more efficient to avoid falling, as each phase can be more flexibly controlled.

Considering that similar coordination patterns are observed in split-belt experiments in both intact and spinal cats (Frigon *et al.*, 2015; Frigon *et al.*, 2017), it is reasonable to assume that drives controlling left and right rhythm generators depend on sensory feedback rather than on supraspinal inputs. One obvious source of sensory feedback is muscle afferent inputs that are known to affect the dynamics of the spinal locomotor CPG circuits (see (Markin *et al.*, 2010) for review). Our model does not explicitly account for this type of feedback. Therefore, the functional interactions and intrinsic flexor and extensor half-centers’ oscillatory properties can be defined in part by inputs from somatosensory afferents. Another type of sensory feedback known to influence locomotion is from the skin (Hurteau *et al.*, 2018). Cutaneous feedback modulation by paw anesthesia alters margins of stability during split-belt cat locomotion (Park *et al.*, 2019). It was recently suggested that this alteration occurs due to misrepresentation of the center of mass in the cat’s balance control system after disrupting cutaneous feedback from the paws (Latash *et al.*, 2020). Altogether, the balance control system (or some of its elements) and locomotor pattern generation may interact at the spinal level, which opens new ways to mathematically model these interactions and thus generate new hypotheses about neuronal pathways mapping somatosensory afferents to the spinal locomotor circuits. Decomposing the functional interactions between left and right RGs into components mediated by local commissural interneurons and spinal reflexes can be a major future research direction where mathematical modeling proves instrumental.

## Acknowledgements

This study was supported by several grants from the National Institutes of Health (R01 NS090919; R01 NS100928; R01 NS110550, R01 NS112304; R01 NS115900), and Discovery Grant from the Natural Sciences and Engineering Research Council of Canada (RGPIN-2016-03790).

